# Low frequency ultrasound enhances chemotherapy sensitivity and induces autophagy in human paclitaxel resistance prostate cancer PC-3R cells through the ERs-mediated PI3K/Akt/mTOR signaling pathway

**DOI:** 10.1101/326629

**Authors:** Yuqi Wu, Xiaobing Liu, Zizhen Qin, Xiangwei Wang

## Abstract

Sonodynamic therapy (SDT) as an emerging tumor-assisting method has attracted a great deal of interest in tumor therapy research in recent years. However, autophagy has been observed in several cancer cells treated with SDT and its role and mechanism are not clear. In the present study, we have investigated the effect of low frequency ultrasound on paclitaxel(PTX) resistance prostate cancer PC-3R cells and demonstrated that low frequency ultrasound could induce cancer cell apoptosis, inhibit the expression of multiple drug resistance genes (MRP3, MRP7 and P-glycoprotein) and reverse drug resistance; we further found that low frequency ultrasound induced autophagy in PC-3R cells. Our results demonstrated that low frequency ultrasound enhanced chemotherapy sensitivity and induced autophagy in PC-3R cells by inhibiting the PI3K/AKT/mTOR pathway. Moreover, we observed that low frequency ultrasound-induced cell autophagy was correlated with endoplasmic reticulum stress (ERs). 4-phenylbutyric acid (4-PBA) - mediated protection against ERs clearly implicated ERs in the activation of autophagy and cell apoptosis. In addition, the results showed that ERs served as an upstream effector of the PI3K/AKT/mTOR pathway. More importantly, we observed that inhibition of low frequency ultrasound-induced autophagy enhanced ERs and improved the role of low frequency ultrasound in reversing drug resistance. Over all, our findings provide new insights into the molecular mechanisms underlying low frequency ultrasound-mediated reversal of drug resistance and autophagy in PC-3R cells and support autophagy as a potential agent for enhancing anti-cancer effect of SDT.

## INTRODUCTION

Prostate cancer is the most common cancer in middle-aged and aged men. It has now become the second major cause of cancer-related death in men ^[1]^. Early prostate cancer is mainly treated with radical surgery, freezing and radiation therapy. For patients with advanced prostate cancer, paclitaxel-based chemotherapy is commonly used after the failure of androgen deprivation therapy. However, there will be drug resistance after a period of treatment, which fails to curb prostate cancer development. Therefore, it is extremely necessary to find a new treatment strategy for prostate cancer ^[2]^.

SDT with low-frequency ultrasonic has strong penetrating ability in biological tissues. In particular, the application of focused ultrasonic can focus the sound energy on deep tissues without injury. Besides, it may contribute to the activation of several ultrasonic-sensitive drugs, such as hematoporphyrin, so as to achieve non-invasive eradication of solid tumors ^[3]^. Recent studies have documented that low-frequency ultrasonic combined with chemotherapeutic drugs can enhance chemotherapy sensitivity and reverse drug resistance in tumor cells ^[4]^.

It has been reported that autophagy can be observed in tumor cells during the application of low-frequency ultrasonic to irradiate cells of nasopharyngeal carcinoma and prostate cancer ^[5,6]^. Nevertheless, its role and mechanism has still been uncleared so far. Autophagy is an evolutionarily conserved process. Autophagosomes perform the recovery of amino acid and energy by encapsulating cytoplasm and organelles, as well as degrading them in lysosomes. The role it plays in the survival and death of cancer cells has always been controversial. Extensive studies have shown that autophagy serves as a protective mechanism in cancer. It can protect cancer cells from various stimuli such as amino acid deficiency, hypoxia, DNA and mitochondrial damage, oxidative stress, etc. ^[7]^. However, it has also been reported that autophagy can inhibit the proliferation of tumor cells and induce cell death (type II programmed cell death) through cooperation with apoptosis ^[8]^. Therefore, exploring the role of autophagy in low-frequency ultrasonic-assisted chemotherapy is quite indispensable to understand the mechanism of reversing drug resistance based on low-frequency ultrasonic, the investigation of which may help to provide new targets and ideas for clinical treatment of reversing drug resistance in prostate cancer.

## MATERIALS AND METHODS

### Cell culture and ultrasound treatment

The PC-3R cell line and Ultrasound treatment instrument (Metron, AA170 type) were provided by the Third Military Medical University. The cells were incubated in RPMI-1640 medium (Gibco) supplemented with 10% fetal bovine serum (FBS; Gibco) and then cultured in a 5% CO2 incubator with saturated humidity at 37°C. Low-frequency ultrasound probe using degassed sterile water as a coupling agent radiated at the bottom of a six-well plate containing 2 mL cell suspension (5×105 cells/ mL). In this study, PC-3R cells were exposed to continuous ultrasound with a frequency of 1MHz, and the spatial average intensity was set to 1.2 W/cm2.

### Cell grouping

PC-3R cells in logarithmic phase were divided into the control group (without any treatment), PTX group (treated with 91.44nM of paclitaxel;Solarbio),Ultrasound group(treated with low-frequency ultrasound),Ultrasound+PTX group(treated with low-frequency ultrasound plus 91.44nM of paclitaxel), Ultrasound+PTX-Atg5 siRNA group (treated with low-frequency ultrasound plus 91.44nM of paclitaxel after transfection of Atg5 siRNA lentivirus;GenePharma), Ultrasound+ 4-PBA group(treated with low-frequency ultrasound plus 10mM of 4-Phenylbutyric acid;Sigma),Ultrasound+ DMSO group(treated with low-frequency ultrasound plus 10mM of Dimethyl sulfoxide, DMSO ; Sigma)

### Ultrasonic cytotoxicity

PC-3R cells were treated with different times of low frequency ultrasound (0, 10, 20, 30, 40,50,60 s). After the treated cells (5×10^5^cells/well) were incubated in a 96-well plate at 37 °C for 24 h, the cell culture medium was removed.CCK-8 (10 μL, Dojindo) solution dissolved in the 100μL medium was added into each well at dark for incubation for 1 h. Then the optical density (OD) of each well at a 450 nm wavelength was detected by microplate reader. The percentage of cytotoxicity was calculated using the following equation: Cytotoxicity (%) = (OD control group - OD ultrasound group)/OD control group × 100%.

### Transmission electron microscopy

Transmission electron microscopy (TEM) was performed to identify mitochondrial morphological changes and autophagy of the PC-3R cells 24 h after ultrasound exposure. Fixed cells were post-fixed in 2% OsO4, dehydrated in graded alcohol and flat embedded in Epon 812 (Electron Microscopy Sciences, Fort Washington, PA, USA). Ultra-thin sections (100 nm) were prepared, stained with uranyl acetate and lead citrate, and examined under an electron microscopy (H-600; Hitachi, Japan).

### Western blotting

Cells were lysed in RIPA buffer (1% NP-40, 0.5% sodium deoxycholate, 0.1% SDS in PBS) containing a Complete Protease Inhibitor Cocktail (Roche, Indianapolis, IN, USA). Protein concentration was determined by Bio-Rad DC protein assay (Bio-Rad, Hercules, CA, USA). Total protein (30 μg) from the cell lysate was separated by SDS-PAGE and transferred to a nitrocellulose membrane. The membrane was blocked in 5% non-fat milk in PBS overnight and then incubated with primary antibody at 37°C for 2 h and washed by tris-buffered saline tween-20 (TBST). Primary antibodies were added, the mixture was incubated overnight. The incubated mixture was washed with TBST, and horseradish peroxidase (HRP) secondary antibodies were added, and the mixture was incubated 37°C for 1 h. The incubated mixture was washed with TBST. HRP electrogenerated chemiluminescence (ECL) was used to develop, the developed films were taken and rinsed with pure water, and the washed films were dried. Scanning was used for recording. The anti-GRP78 was used at a 1:1000 dilution (Abcam), the anti-PI3 Kinase p110 beta was used at a 1:600 dilution (Abcam), the anti-AKT1 (phospho S473) was used at a 1:1000 dilution (Abcam), the anti-mTOR (phospho S2481) was used at a 1:1000 dilution (Abcam), the anti-LC3B was used at a 1:1000 dilution (Abcam), the anti-APG5L/ATG5 was used at a 1:1000 dilution (Abcam), the anti-NF-kB p65 was used at a 1:1000 dilution (Abcam), the anti-Bcl-2 was used at a 1:1000 dilution (Abcam) and the anti-beta Actin protein antibody was used at a 1:200 (Abcam).

### Apoptosis assay

Apoptosis was detected using an Annexin V-PE Apoptosis Detection Kit (BD Pharmingen TM, San Jose, CA, USA). Cells in the logarithmic growth phase were harvested and washed twice in PBS. Then, 1×10^6^ cells were counted and washed twice in PBS before re-suspension in 1 X Binding Buffer. PE Annexin V (5 ul) and 7-AAD (5 ul) were added to the cells on ice for 30 min to permit cell staining, followed by the addition of 400 ul 1 X Binding Buffer to each sample.

### Cell proliferation assay

Cell proliferation was evaluated using a Cell-Counting Kit 8 (CCK-8), according to the manufacturer’s instructions (Donjindo). After different treatment for 24 h, PC-3R cells were seeded in 96-well plates (2,000 cells/well) and 10 ul CCK-8 solution was added to each well at the indicated time point after transfection. The cells were then incubated for a further 2 h. The absorbance of the cultures was then measured at 450 nm on a Multicacan FC Microplate Photometer (Thermo Fisher Scientic, Rochester, NY, USA).

### Isolation of total RNA and qRT-PCR

Total RNA was extracted from harvested cells with Donjindo (Donjindo Molecular Technologies, Inc., Kumannoto, Japan), according to the manufacturer’s instructions and quantitative reverse transcription PCR (qRT-PCR) was conducted as previously described. Briefly, for the detection of Bcl-2, MRP3, MRP7 and P-glycoprotein, 1 ug total RNA per sample was converted to cDNA using a cDNA synthesis Kit (Invitrogen, Carlsbad, CA, USA). These cDNAs were amplified and detected using a SYBR Green PCR kit (Qiagen, Valencia, CA, USA). GAPDH was used as an endogenous control. For detection of the mRNA, cDNA products were synthesized using miScript, Reverse Transcription Kit (Qiagen). Primers specific for Bcl-2, MRP3, MRP7, P-glycoprotein or endogenous control U6 were purchased from Qiagen (Table 1). qRT-PCR was performed using miScript SYBR Green PCR Kit (Qiagen). All reactions were run in triplicate on a Bio-Rad C1000 thermal cycler (CFX-96 real-time PCR detection systems, Bio-Rad). Fold changes in mRNA expression was calculated according to the 2–ΔΔCT method.

**TABLES.**
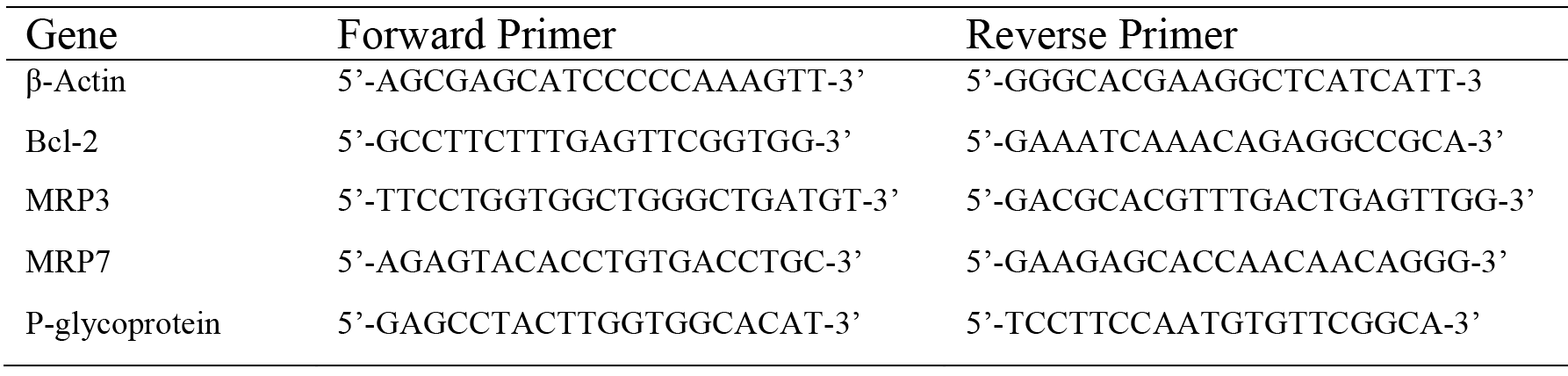

## RESULTS

### Cytotoxicity of ultrasound in PC-3R cells

To assess the non-cytotoxicity exposure time of ultrasound in the PC-3R cell line, the ultrasound-treated cells were incubated for 24 h. The cytotoxic effect of ultrasound on the PC-3R cells is shown in Fig. 1. The death rate increased along with the ultrasound exposure time, showing that ultrasound treatment definitively killed PC-3R cells. In order to avoid the interference of ultrasonic cytotoxicity on the experiment we choose 10s exposure time (cytotoxicity less than 10%) for subsequent experiments.

**Figure 1.**
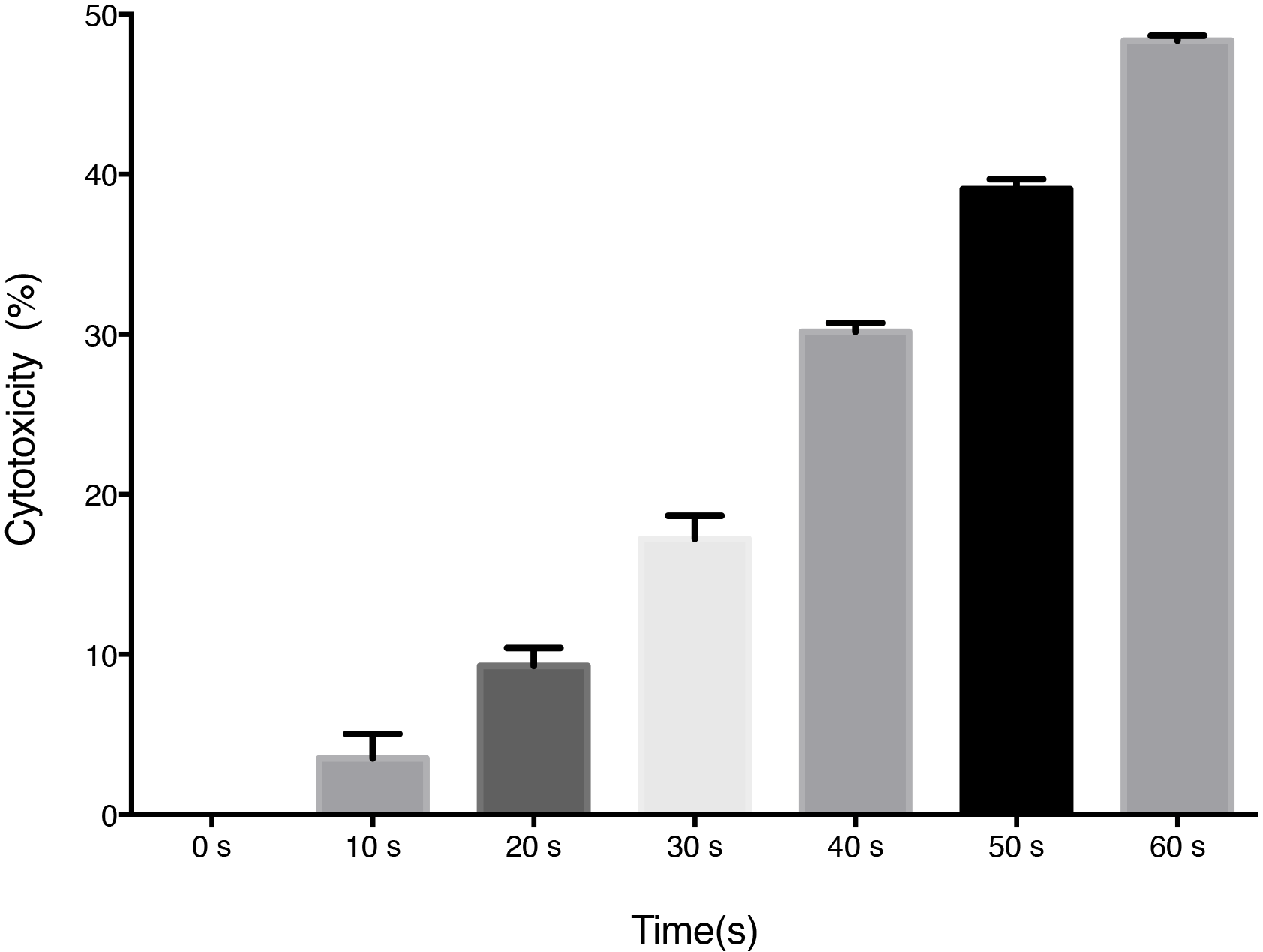
Cytotoxic effect of ultrasound on PC-3R cells. PC-3R cells cultured for 24h was assessed by CCK-8 assay after ultrasound exposure. The frequency of ultrasound was 1 MHz and the spatial average ultrasonic intensity was set at 1.2 W/cm2.

### Low-frequency ultrasound induces autophagy in PC-3R cells

To examine whether ultrasound induced autophagy in PC-3R cells, several autophagy assays were performed. First, autophagy was investigated using TEM. As shown in Fig. 2A, swollen mitochondria, vacuoles and autophagosomes were noted in the cells treated by ultrasound. Moreover, treatment with ultrasound combined with paclitaxel, autophagosomes increased significantly in the PC-3R cells. However, pretreatment of cells with Atg5 siRNA number of autophagosome reduced after treated by ultrasound combined with paclitaxel.

**Figure 2.**
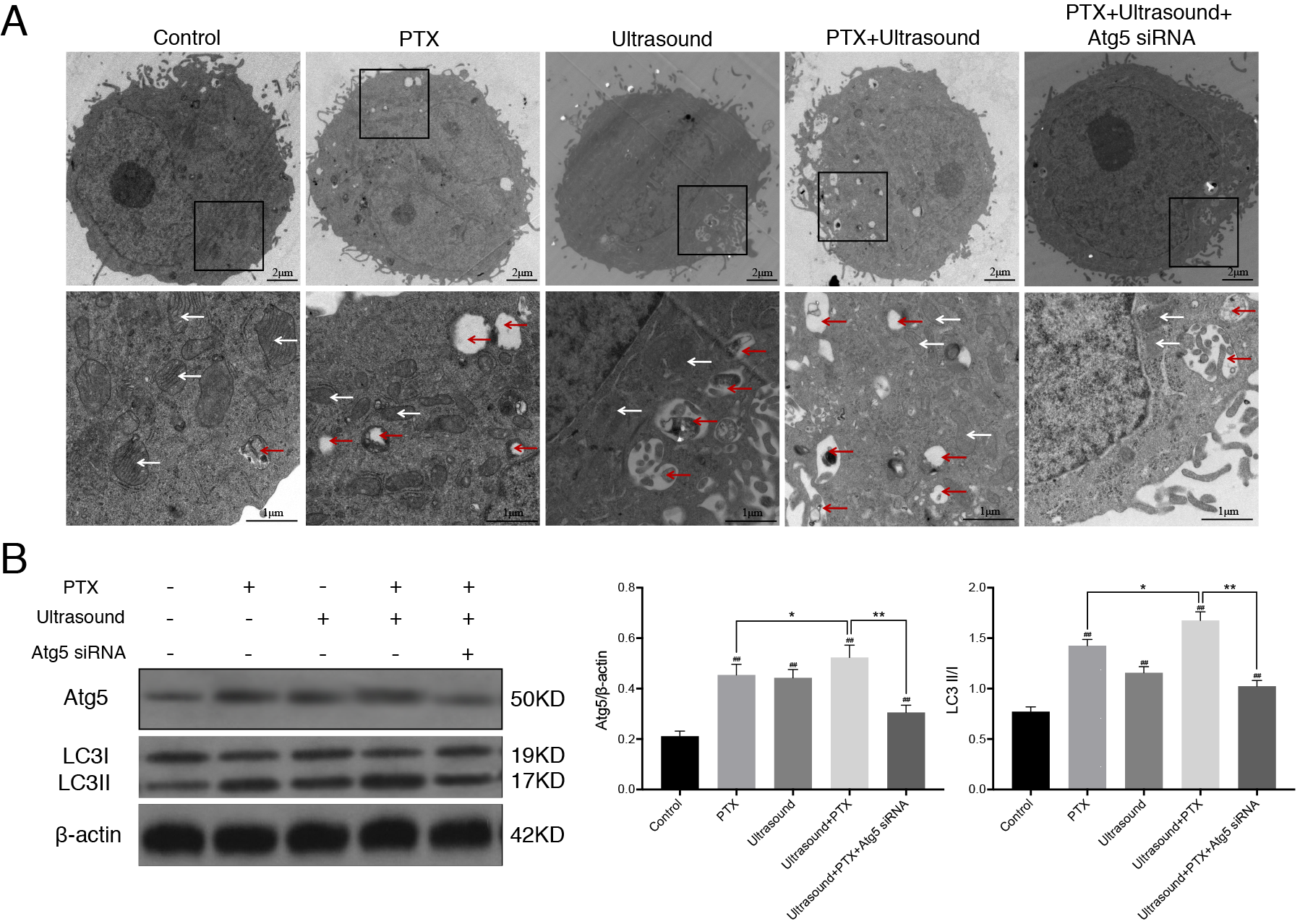
The effect of ultrasound on autophagy in PC-3R cells. (A) Investigation of ultrastructural morphology and autophagy of PC-3R cells was used by TEM at 24 h after ultrasound treatment (10 s) combined with or without paclitaxel(91.44nM) or Atg5 siRNA. The red arrow indicates the autophagosome and the white arrow indicates the mitochondria. (B) Cells were untreated or treated with ultrasound (10 s) in the presence or absence of Atg5 siRNA or paclitaxel(91.44nM) for 24 h. The expression levels of Atg5 and LC3B were assessed. Protein lysates were collected and assayed by western blotting. *p<0.05, **p<0.01, ^##^p<0.01, as compared with control group.

Furthermore, we also examined the effect of low-frequency ultrasound on level of autophagy using a western blot analysis. As shown in Fig. 2B, the protein expression levels of Atg5 and LC3 II were significantly increased after treated by ultrasound or cotreatment with ultrasound and paclitaxel. Conversely, these two protein level was obviously decreased after using Atg5 siRNA to block autophagy in cells. These results showed that low-frequency ultrasound and paclitaxel induces autophagy in PC-3R cells and Atg5 siRNA inhibits effectively the autophagy.

### Low-frequency ultrasound induces apoptosis and enhances chemotherapy sensitivity in PC-3R cells

As shown in Fig. 3A, flow cytometry was used to quantify the apoptotic rate of prostate cancer cell lines. Here, we found that the apoptotic rate of PC-3R cells treatment with ultrasound combined with paclitaxel was higher than that of PC-3R cells treatment with paclitaxel only, and the apoptotic rate of PC-3R cells transfected with Atg5 siRNA was significantly increased compared with cells treatment with ultrasound and paclitaxel (P<0.01).

**Figure 3.**
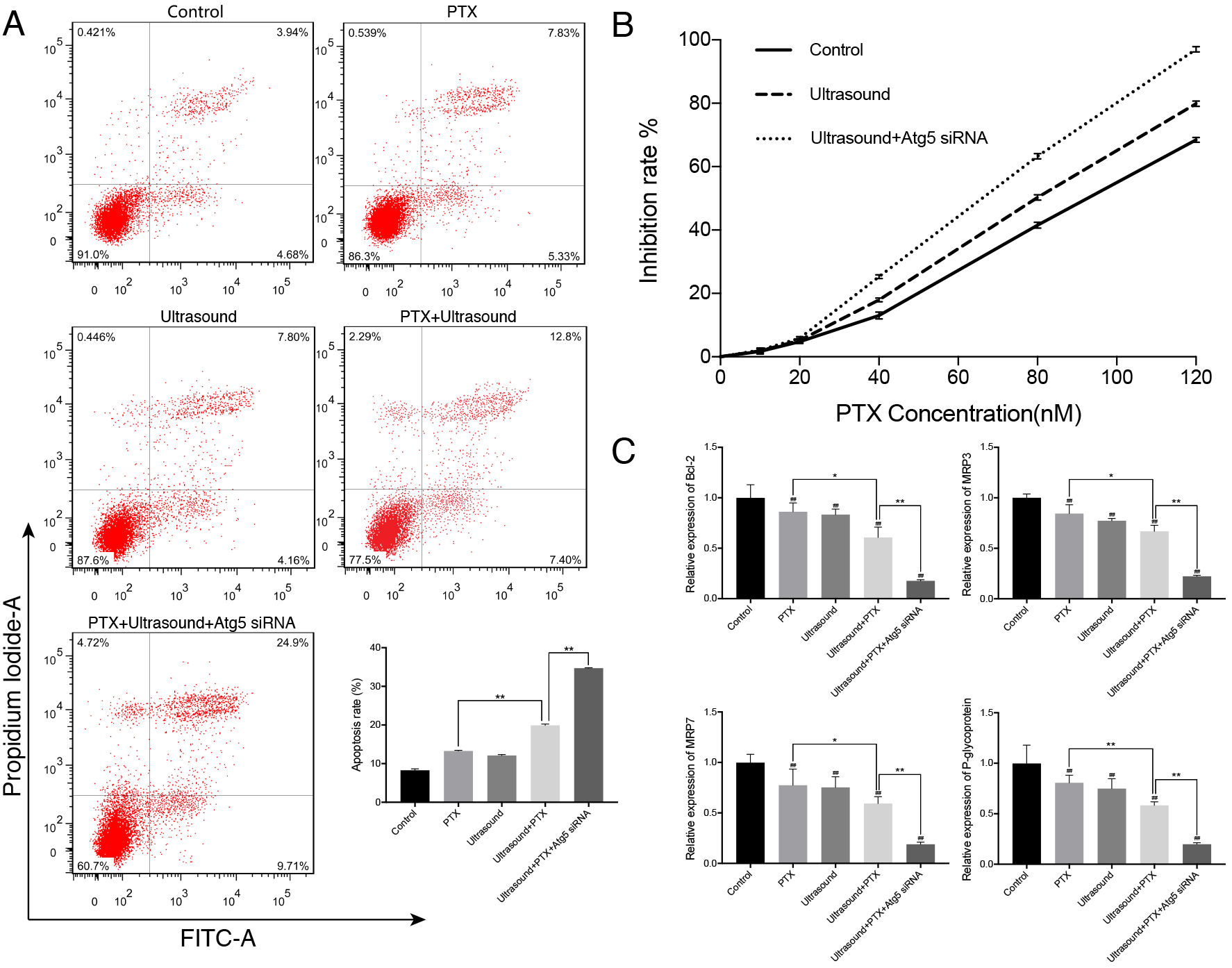
The effect of ultrasound on apoptosis and chemotherapy sensitivity in PC-3R cells. PC-3R cells were untreated or treated with ultrasound (10 s) in the presence or absence of Atg5 siRNA or paclitaxel(91.44nM) for 24 h, (A)stained with Annexin V-FITC and PI, then measured by flow cytometry. (C)The expression levels of Bcl-2, MPR3, MPR7 and P-glycoprotein were assessed by qPCR. *p<0.05, **p<0.01, ^##^p<0.01, as compared with control group. (B)PC-3R cells treated with ultrasound (10 s) in the presence or absence of Atg5 siRNA were treated with different concentrations of paclitaxel for 24h. The cell inhibition was assessed by CCK-8 assay.

Furthermore, to explore the effect of ultrasound on chemotherapy sensitivity in PC-3R cells CCK-8 assay was used. We observed that there was a significant dose-dependent ascent in paclitaxel inhibition rate of prostate cancer cells. Moreover, paclitaxel inhibition rate of PC-3R cells treatment with ultrasound was higher than control group and it lower than PC-3R cells treatment with Atg5 siRNA and low-frequency ultrasound (Fig. 3B). Moreover, we analyzed the gene expression levels of the molecules that are known to be involved in the apoptosis and drug resistance. As shown in Fig.3C, the expression level of Bcl-2, MPR3, MPR7 and P-glycoprotein were decreased after treated by ultrasound and paclitaxel. Importantly, the expression level of these genes was significantly decreased in PC-3R cells transfected with Atg5 siRNA compared with cells co-treatment with ultrasound and paclitaxel. These findings strongly suggest that low-frequency ultrasound contributed to enhancement paclitaxel sensitivity by inducing apoptosis and inhibiting resistance-related genes in PC-3R cells and ultrasound-induced autophagy played the role of cytoprotector in the process of low frequency ultrasound reversal of drug resistance.

### Low-frequency ultrasound induces apoptosis through the ER-mediated PI3K/AKT/mTOR signaling pathway in PC-3R cells

To examine the effect of low-frequency ultrasound on endoplasmic reticulum stress, we used a western blot analysis for GRP78, a ER marker protein. As shown in Fig. 4, the protein expression levels of GRP78 were significantly increased after treated by ultrasound or co-treatment with ultrasound and paclitaxel. Furthermore, in regard to the interplay between endoplasmic reticulum stress and apoptosis, we further elucidated the underlying molecular mechanism. As shown in Fig. 4, the protein expression levels of PI3K, p-AKT, mTORC1, Bcl-2 and p65(NF-kB related protein) were significantly decreased after treatment with ultrasound and further decreased after co-treatment with ultrasound and paclitaxel. Interestingly, we found that the protein expression levels of GRP78 were significantly increase strongly after treatment with Atg5 siRNA, while the PI3K, p-AKT, mTORC1, Bcl-2 and p65 protein level was obviously decreased. Taken together, these changes in the expression levels of PI3K/AKT/mTOR signaling pathway-related proteins indicated that ultrasound induces apoptosis through ER-mediated PI3K/AKT/mTOR signaling pathway in PC-3R cells. At the same time, inhibition of autophagy might enhance signaling pathway.

**Figure 4.**
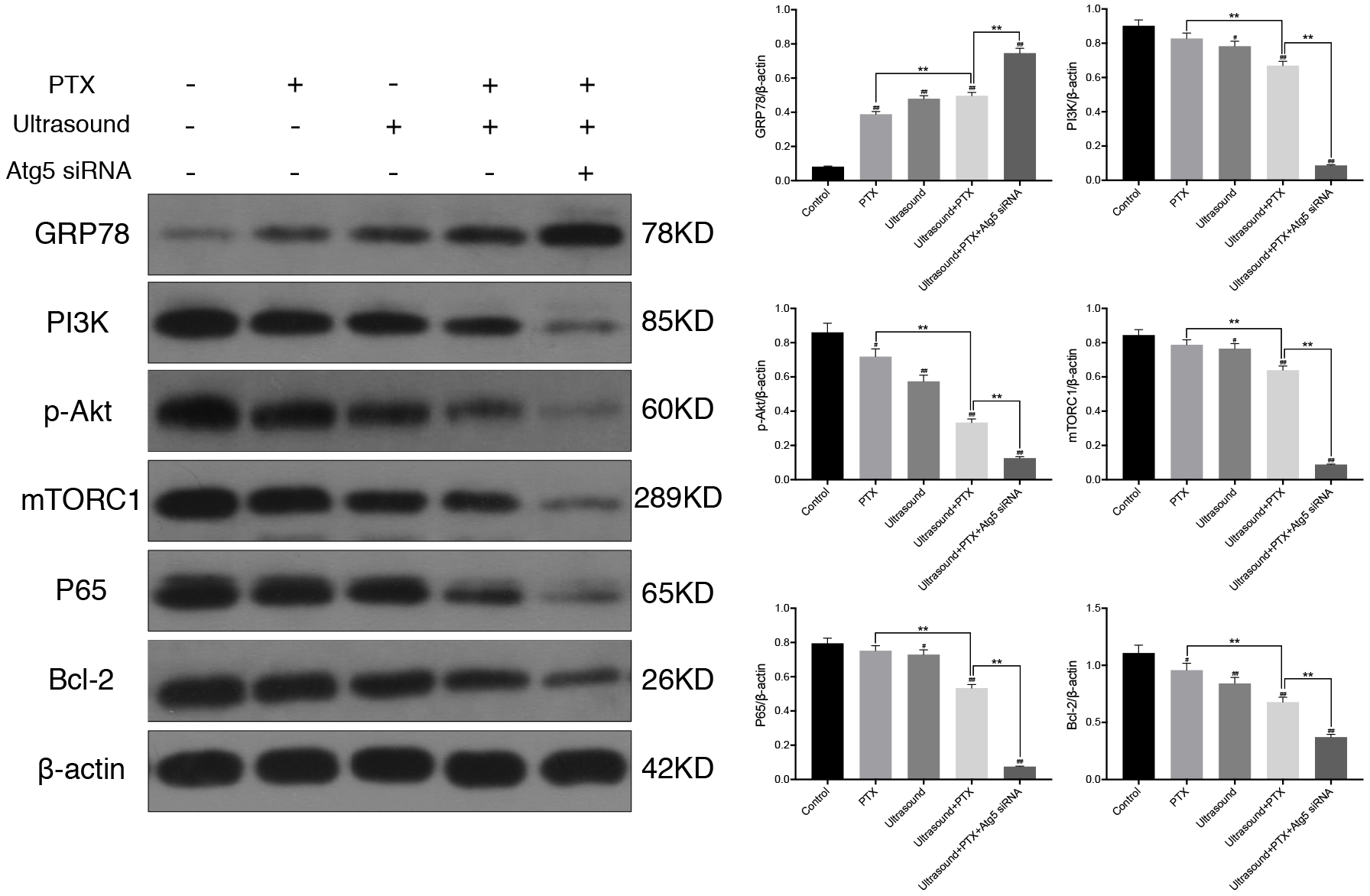
The effect of ultrasound on ERs-mediated PI3K/Akt/mTOR signaling pathway in PC-3R cells. PC-3R cells were untreated or treated with ultrasound (10 s) in the presence or absence of Atg5 siRNA or paclitaxel(91.44nM) for 24 h. The expression levels of PI3K, p-AKT, mTORC1, Bcl-2 and p65 were assessed. Protein lysates were collected and assayed by western blotting. *p<0.05, **p<0.01, ^#^p<0.05, ^##^p<0.01, as compared with control group.

### Low-frequency ultrasound induces autophagy through the ER-mediated PI3K/AKT/mTOR signaling pathway in PC-3R cells

To examine the relation between endoplasmic reticulum stress and autophagy, we treated PC-3R cell with 4-PBA, an endoplasmic reticulum stress inhibitor. Autophagy was investigated using TEM. As shown in Fig. 5A, autophagosomes increased significantly in the PC-3R cells treatment with ultrasound. However, the number of autophagosome reduced after treatment with 10mM 4-PBA. Furthermore, we also used immunofluorescence staining to detect autophagy. As shown in Fig. 5B, green fluorescence was emitted in cells treated with ultrasound. However, treatment of cells with 4-PBA decreased the green fluorescence intensity after treatment with ultrasound. In addition, the interplay between endoplasmic reticulum stress and autophagy was further studied by assessing the levels of ER-mediated PI3K/AKT/mTOR signaling pathway-associated proteins. As shown in Fig.5C, a significant decrease in the expression of GRP78 and LC3II proteins was observed, while expression of PI3K, p-AKT and mTORC1 increased after treatment with 4-PBA and ultrasound. Consistent with these results, we found that ultrasound induces autophagy through ER-mediated PI3K/AKT/mTOR signaling pathway in PC-3R cells.

**Figure 5.**
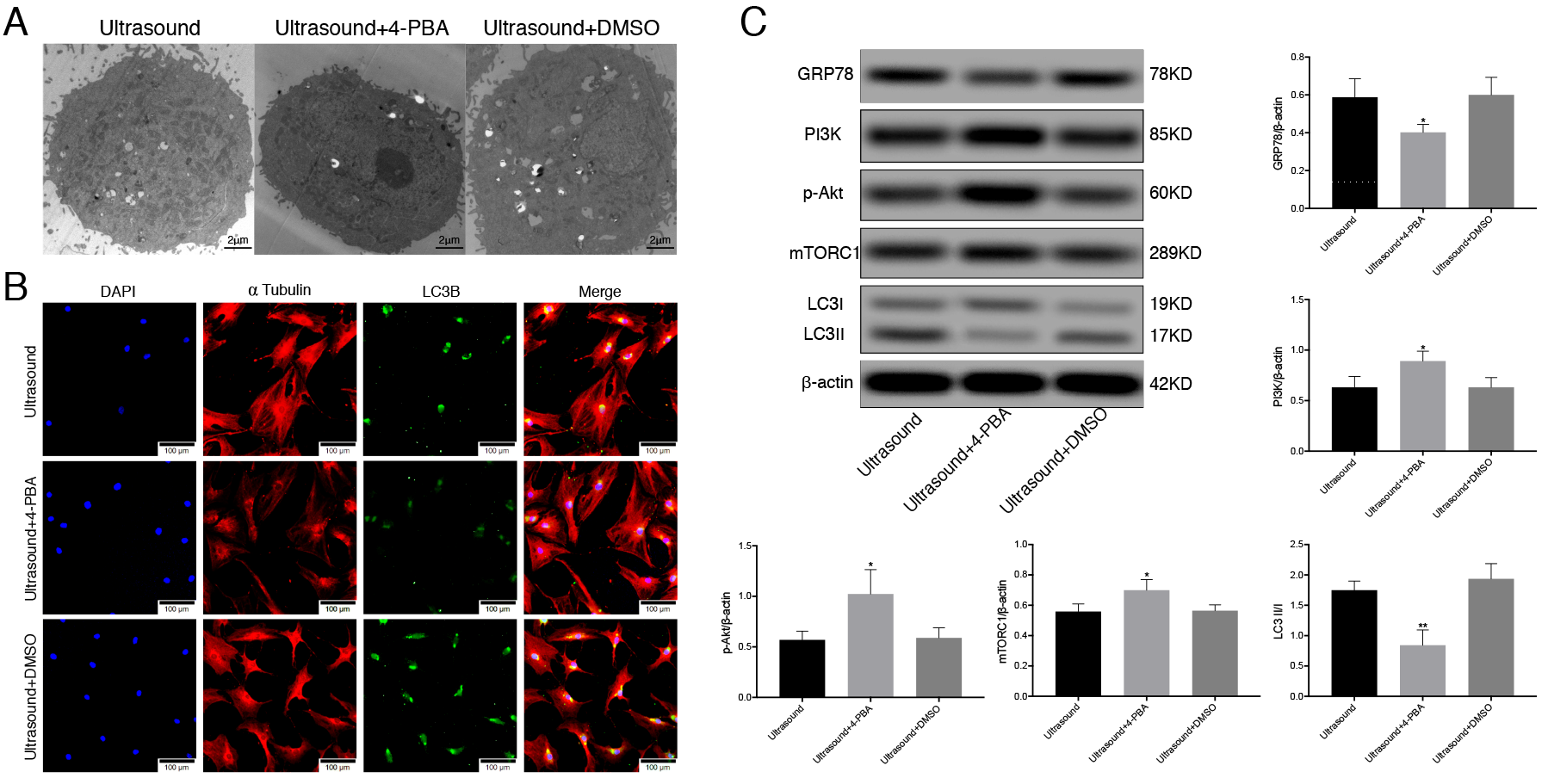
Low-frequency ultrasound induces autophagy through the ER-mediated PI3K/AKT/mTOR signaling pathway in PC-3R cells. PC-3R cells were treated with ultrasound (10 s) in the presence or absence of 4-PBA(10mM) or DMSO(10mM) for 24 h. (A)Investigation of ultrastructural morphology and autophagy was used by TEM and (B)the level of autophagy was detected by immunofluorescence staining. The expression levels of PI3K, p-AKT, mTORC1 and LC3B were assessed. Protein lysates were collected and assayed by western blotting. *p<0.05, **p<0.01, as compared with ultrasound group.

## DISCUSSION

Studies have shown that low-frequency ultrasonic can selectively increase the permeability of tumor cell membranes in breast cancer, ovarian cancer and neuroblastoma. Simultaneously, it can activate mitochondria-caspase signaling pathway and down-regulate the expression of ATP-binding cassette(ABC) transporter, so as to improve the sensitivity of tumors to chemotherapy drugs and reverse drug resistance ^[4]^. The present study examined the drug resistance reversing effect of low-frequency ultrasonic in paclitaxel-resistant prostate cancer cells. Corresponding results revealed that autophagy induced by low-frequency ultrasonic could be enhanced when the latter was combined with paclitaxel on prostate cancer cells. By using Atg5 siRNA to inhibit autophagy, the anti-apoptotic gene Bcl-2 was observed to be further down-regulated with a significant increase of the apoptosis rate. MRP3, MRP7 and P-glycoprotein are members of the ABC transporter family, by which chemotherapeutic drugs can be pumped to reduce the sensitivity of paclitaxel to cells. After the inhibition of autophagy, their transcription level was reduced, accompanied by increased concentration of intracellular chemotherapeutic drugs, resulting in significant enhancement of the efficiency of low-frequency ultrasonic in reversing drug resistance. The above suggests that autophagy can exert a protective role for cells during low-frequency ultrasonic-assisted paclitaxel treatment for prostate cancer.

Endoplasmic reticulum, as the central organelle in cells, is involved in the secretory function, as well as the folding, transfer and post-translocational modification of secretory protein. Dysfunction of the endoplasmic reticulum would lead to the accumulation of misfolded and unfolded proteins, the process of which is called Endoplasmic Reticulum Stress. A large number of prior studies have shown that endoplasmic reticulum stress may be induced by chemotherapeutic drugs ^[9]^. However, low-frequency ultrasonic has rarely been reported to cause endoplasmic reticulum stress. Our results indicated that endoplasmic reticulum stress occurred in prostate cancer cells after low-frequency ultrasonic irradiation at non-toxic time, which may be related to the low-frequency ultrasonic cavitation. When ultrasonic is applied to cancer cells, bubbles will oscillate in cells, leading to the flow of the surrounding liquid, which results in the mixture of the surrounding cytoplasm. Meanwhile, bubbles will continue to enlarge until implosion occurs ^[10]^. The function of the endoplasmic reticulum can eventually be affected by this process, resulting in endoplasmic reticulum stress. In the treatment of pancreatic cancer with pentafluoride, Alok Ranjan et al. found that pentafluride could induce autophagy by triggering endoplasmic reticulum stress ^[11]^. Furthermore, the autophagy level decreased after the blocking of endoplasmic reticulum stress by endoplasmic reticulum stress blockers or CHOP siRNA. Our experimental results also revealed that endoplasmic reticulum stress would occur and the level of autophagy would increase with low-frequency ultrasonic irradiation of cells. When low-frequency ultrasonic was combined with paclitaxel, endoplasmic reticulum stress was enhanced and the autophagy level increased correspondingly. However, a decline of ultrasonically induced autophagy was exhibited when cells were treated with endoplasmic reticulum stress inhibitor of 4-phenylbutyrate. Interestingly, it was also discovered in our study that endoplasmic reticulum stress increased obviously when autophagy was inhibited. It is possibly due to the fact that autophagy can provide energy and materials in the process of “self-cannibalization” at the same time when endoplasmic reticulum stress induces autophagy to improve intracellular environment, thereby inhibiting the occurrence of endoplasmic reticulum stress. In addition, there are also other studies showing that autophagy can remove reactive oxygen species (ROS) and misfolded proteins to avoid their accumulation in cells. Among them, the accumulation of ROS and protein are considered important factors in inducing endoplasmic reticulum stress ^[12]^.

Yang N et al. carried out an experiment focusing on the treatment of ovarian cancer cells with tunicamycin. Their results revealed that endoplasmic reticulum stress could inhibit the PI3K/AKT/mTOR signaling pathway to induce apoptosis in human ovarian cancer cells ^[13]^. Crucially, PI3K/AKT/mTOR is a classical signaling pathway involved in the regulation of autophagy. The activation of class I PI3 kinase (PI3K) can phosphorylate ptdIns(4,5)P2 and form PtdIns(3,4,5)P3 in the plasma membrane. In the meantime, class I PI3 kinase (PI3K) can also recruit protein kinase B and phosphoinositide-dependent protein kinase 1, and the co-recruitment of these two kinases leads to the activation of AKT. Subsequently, via the phosphorylation of TSC1, the accumulation of RHEB GTP can be promoted by active AKT, while the former is capable of activating the mammalian target of rapamycin (mTOR) ^[14]^. Furthermore, mTOR can activate its downstream ribosomal protein S6 (p7s6), and the phosphorylation of S6 (p7s6) can inhibit the occurrence of autophagy ^[15]^. As is revealed in our experimental results, PI3K/AKT/mTOR pathway was also inhibited when low-frequency ultrasonic treatment was applied to induce endoplasmic reticulum stress. Meanwhile, in the combined use of low-frequency ultrasonic and paclitaxel, PI3K, p-Akt and mTORC1 were further down-regulated with the enhancement of endoplasmic reticulum stress. Following the application of ATG5 siRNA to inhibit autophagy, endoplasmic reticulum stress was further enhanced, which in turn resulted in significant down-regulation of PI3K/AKT/mTOR pathway. After the inhibition of endoplasmic reticulum stress, the inhibitory effect of ultrasonic on PI3K/AKT/mTOR pathway was weakened correspondingly. Therefore, it is assumed that low-frequency ultrasonic induces autophagy by triggering endoplasmic reticulum stress which inhibits PI3K/AKT/mTOR signaling pathway. When low-frequency ultrasonic is combined with paclitaxel, the endoplasmic reticulum stress will be enhanced along with the down-regulation of PI3K/AKT/mTOR pathway and the rise of autophagy levels. By contrast, with autophagy inhibited, the regulative effect of autophagy can be reduced on the intracellular environment, accompanied by further enhancement in endoplasmic reticulum stress, thereby leading to the increased inhibitory effect on PI3K/AKT/mTOR pathway. mTOR is the main regulator for cell-growth control which participate in drug resistance of tumor cells and can inhibit tumor apoptosis by regulating the expression of NF-κB and ABC transporters ^[16]^. As is illustrated by our experimental results, expression levels of P65, MRP3, MRP7 and P-glycoprotein were down-regulated correspondingly with the decrease of mTORC1 expression. On the other hand, PI3K/Akt signal transduction pathway is regarded as the chief pathway for the survival of tumor cells. AKT can phosphorylate a number of its downstream substrates to facilitate the growth and proliferation of tumor cells, inhibit cell apoptosis, and enhance tumor cell resistance to chemotherapy and radiotherapy involving Bcl-2 family and apoptosis inhibitory protein family, etc. In addition, the transcriptional activity of NF-κB can be strengthened by phosphorylating eNOS ^[17,18]^, as was proved by our experimental results.

To sum up, the cavitation of low-frequency ultrasonic can trigger endoplasmic reticulum stress which down-regulates PI3K/AKT/mTOR signal transduction pathway to reduce the transcriptional activity of ABC transporter, Bcl-2 and NF-κB, so as to increase intracellular concentration of paclitaxel, promote apoptosis and reverse drug resistance. On the other hand, PI3K/AKT/mTOR signal transduction pathway down-regulated by endoplasmic reticulum stress can induce autophagy of paclitaxel-resistant cells in prostate cancer in the process of reversing tumor cells drug resistance by low-frequency ultrasonic, while autophagy may, in turn, inhibit endoplasmic reticulum stress by eliminating ROS and misfolded proteins. As a result of autophagy inhibition, endoplasmic reticulum stress is enhanced, inducing further down-regulation of PI3K/AKT/mTOR signal transduction pathway. Consequently, a further drop in the transcriptional activity of ABC transporter, Bcl-2 and NF-κB can be observed together with an increase of intracellular concentration of paclitaxel and an enhancement of cell apoptosis. Hence, the efficiency in reversal of drug resistance is significantly improved by low-frequency ultrasonic.

## ACKNOWLEDGEMENTS

The study was sponsored by grant 81172443 from the National Natural Science Foundation of China. No benefits in any form have been or will be received from a commercial party related directly or indirectly to the subject of this study.

